# Antibiotic-resistance mutations in penicillin-binding protein 2 from the ceftriaxone-resistant *Neisseria gonorrhoeae* strain H041 strike a delicate balance between increasing resistance and maintaining transpeptidase activity

**DOI:** 10.1101/2025.11.13.688206

**Authors:** Marissa M. Bivins, Joshua Tomberg, Madeleine Bagshaw, Avinash Singh, Sandeepchowdary Bala, Christopher Davies, Robert A. Nicholas

## Abstract

The mosaic *penA* allele (*penA41*) from H041, the most ceftriaxone-resistant *Neisseria gonorrhoeae* strain identified to date, encodes a variant of the essential Penicillin-Binding Protein 2 (PBP2) with 60 amino acid mutations compared to PBP2 from the antimicrobial-susceptible strain, FA19. Based on previous work from our lab and others, we identified a minimal set of 10 mutations that, when introduced into the β-lactam antibiotic-susceptible *penA* allele from FA19 (*penA19*), confers two-thirds of the ceftriaxone and cefixime resistance compared to the *penA41* allele. Three mutations (A311V, I312M, and V316P) are in the 𝛼2 helix of PBP2 containing the catalytic serine (Ser310), two (F504L and N512Y) are in the 𝛽3-𝛽4 loop that is important in binding and acylation, and one (G545S) interacts with conserved amino acids in the active site. The seventh mutation, T483S, confers substantial resistance to ceftriaxone within the minimal mutant set but requires the presence of three epistatic mutations located on the other side of the protein that do not alter resistance on their own yet are necessary to retain essential transpeptidase activity. These epistatic mutations change the backbone dihedral angles at position-447, which may increase flexibility of the enzyme and help restore essential transpeptidation. Our results highlight the complex balance necessary for evolving cephalosporin resistance while also retaining sufficient transpeptidase function in PBP2.

**Author Summary:** In this study, we set out to understand how Neisseria gonorrhoeae, the bacterium that causes gonorrhea, is able to resist the last remaining recommended antibiotic, ceftriaxone. Gonorrhea is a common sexually transmitted infection worldwide, and rising resistance threatens to make it untreatable. We focused on penicillin-binding protein 2 (PBP2), which is essential for the bacterium’s survival and is the lethal target of ceftriaxone. By incorporating a subset of the 60 PBP2 mutations found in a highly resistant strain into PBP2 from an antibiotic-susceptible strain, we discovered that resistance evolved from a combination of mutations that work together to directly reduce the capacity of ceftriaxone to inactivate the protein and others that act as “supporting” mutations to keep the protein functional despite the presence of the resistance mutations. Our study highlights how *N. gonorrhoeae* successfully negotiates the delicate balance between resistance and function to escape the lethal action of antibiotics.

## Introduction

*Neisseria gonorrhoeae* is a Gram-negative bacterium that causes the sexually transmitted infection (STI), gonorrhea. The World Health Organization (WHO) reported an estimated 82 million gonorrhea cases globally in 2019 [1] and the Centers for Disease Control and Prevention (CDC) reported that gonorrhea is the second most prevalent STI in the United States, with over 600,000 cases reported in 2023 [2]. The actual number of gonorrhea cases is likely to be much higher, since 7% to 15% of infections in men and 16% to 70% of infections in women are asymptomatic [3–5]. Prolonged *N. gonorrhoeae* infections can result in serious complications, including pelvic inflammatory disease, infertility, and increased susceptibility to HIV infection [6–8]. Currently, the only recommended treatment by the CDC for gonorrhea is a single 500 mg injection of the extended spectrum cephalosporin (ESC), ceftriaxone, but resistance to this β-lactam antibiotic is on the rise [9–13].

The targets for β-lactam antibiotics such as ceftriaxone are the penicillin-binding proteins (PBPs), which synthesize and cross-link the peptidoglycan layer that surrounds the bacteria [14, 15]. *N. gonorrhoeae* has two essential PBPs: PBP1 and PBP2 [16, 17]. PBP1 is a class A bifunctional PBP that catalyzes both glycan polymerization and crosslinking of the peptide strands of peptidoglycan, whereas PBP2 is a class B monofunctional PBP that cross-links peptide strands during cell division but requires a partner protein, FtsW, for glycan polymerization [18–20]. Peptidoglycan crosslinking occurs through transpeptidation, in which the PBPs form an acyl-enzyme complex with the penultimate D-Ala from the acyl-D-Ala-D-Ala C-terminus of the peptidyl chain and then reacts with the free amino group in *m*-diaminopimelic acid (*m*-DAP) from a parallel peptidoglycan strand, forming a cross-link and releasing the enzyme to catalyze another round. β-lactam antibiotics target PBPs by mimicking the acyl-D-Ala-D-Ala C-terminus of the peptide chains and forming a long-lived acyl-enzyme complex with the PBP [14, 15, 21, 22].

Of the two essential PBPs, PBP2 historically has had a much higher rate of acylation than PBP1 with essentially all β-lactam antibiotics used to treat gonococcal infections, making PBP2 the lethal target of these antibiotics [16, 23, 24]. During the four decades when penicillin was used to treat infections, the *penA* gene encoding PBP2 acquired mutations comprising an amino acid insertion (Asp345a) and four to eight mutations in the C-terminus of the PBP, with the insertion and the C-terminal mutations contributing equally to decreasing the acylation rate [24, 25]. Following the removal of penicillin, the ESCs ceftriaxone and cefixime were used extensively to treat gonorrhea, but strains emerged from East Asia in the early 2000s with decreased susceptibility to ESCs [26–28]. Unlike penicillin-resistant strains, these isolates had so-called mosaic *penA* alleles, in which regions from the *penA* genes of other *Neisseria* species were recombined into *N. gonorrhoeae penA*, resulting in PBP2 variants with 40-60 mutations compared to PBP2 from susceptible strains [26, 29]. These mosaic alleles emerged as a result of *N. gonorrhoeae* being naturally competent, which facilitates the uptake and recombination of closely related DNA into its genome by homologous recombination. This property allows for both the original formation of the mosaic alleles (a low frequency event) and the spread of these alleles in the population (high frequency), provided that these genes do not have a large fitness cost.

One example of a strain with a mosaic *penA* allele is H041 (referred to as *penA41*), which was isolated in Japan in 2009 and has the highest level of resistance to ceftriaxone (MIC = 2-4 µg/mL) and cefixime (MIC = 8 µg/mL) of any gonococcal isolate to date [30, 31]. Compared to PBP2 from an antibiotic-susceptible isolate, e.g. FA19 (PBP2^FA19^), PBP2 from H041 (PBP2^H041^) has 60 amino acid alterations [30]. To understand how these mutations decrease the capacity of PBP2 from ESC-resistant strains to be inhibited by ESCs, we previously solved crystal structures of the transpeptidase domains of PBP2^FA19^ and PBP2^H041^ (denoted by the prefix t) and found that conformational changes observed in tPBP2^FA19^ after acylation by ceftriaxone and ceftriaxone do not occur in tPBP2^H041^ [32, 33]. These changes include movement of the 𝛽3-𝛽4 loop towards the active site, where it forms contacts with other residues in PBP2 and with the R_1_ moieties of ESCs, and twisting of the 𝛽3 strand, which contains the highly conserved KTG motif [22, 34], towards the active site to aid in formation of the oxyanion hole that stabilizes the tetrahedral transition state during acylation [32]. In contrast to tPBP2^FA19^, the 𝛽3-𝛽4 loop in tPBP2^H041^ is “outbent” and pointed away from the active site in the *apo* form and remains so even after acylation with ceftriaxone or cefixime [33].

In previous studies, we and others have identified seven mutations that are important in conferring resistance: A311V, I312M, V316P, T483S, F504L, N512Y, and G545S (Fig. 1) [35–38]. Understanding how these seven mutations work together to impact both antibiotic resistance and strain fitness is vital for understanding *N. gonorrhoeae* evolution more broadly and for developing new therapeutics. We grouped these seven mutations from the *penA41* allele of H041 into three clusters based on their location in the protein and examined their effects on the MICs of ESCs and penicillin G by either incorporating them into the *penA* allele from FA19 (*penA19*) in the absence of other mutations or reverting them to their *penA19* counterparts within the *penA41* background. Each group of mutations had varied effects depending on the antibiotic and the PBP2 background. We also attempted to incorporate all seven mutations together into *penA19* to create a “minimal mutant” that would confer the maximum level of resistance to FA19 after transformation. However, one of the mutations, T483S, was both a crucial resistance mutation and a disrupter of transpeptidase function, and required the presence of three non-resistance mutations (referred to as epistatic mutations) to maintain sufficient transpeptidase activity to support growth. When the seven mutations and three epistatic mutations were incorporated into the *penA19* background and transformed into FA19, they conferred 67% of the MICs of both ceftriaxone and cefixime relative to FA19 *penA41*. Crystal structures of the mutants reveal that the epistatic mutations change the backbone dihedral angles at position 447, which may impart increased flexibility to the enzyme containing the resistance mutations. These data provide insight into how PBP-mediated resistance arises and how resistance mutations combine to increase resistance, and additionally highlight the delicate balance between increasing resistance and preserving essential transpeptidase activity.

**Figure 1.**
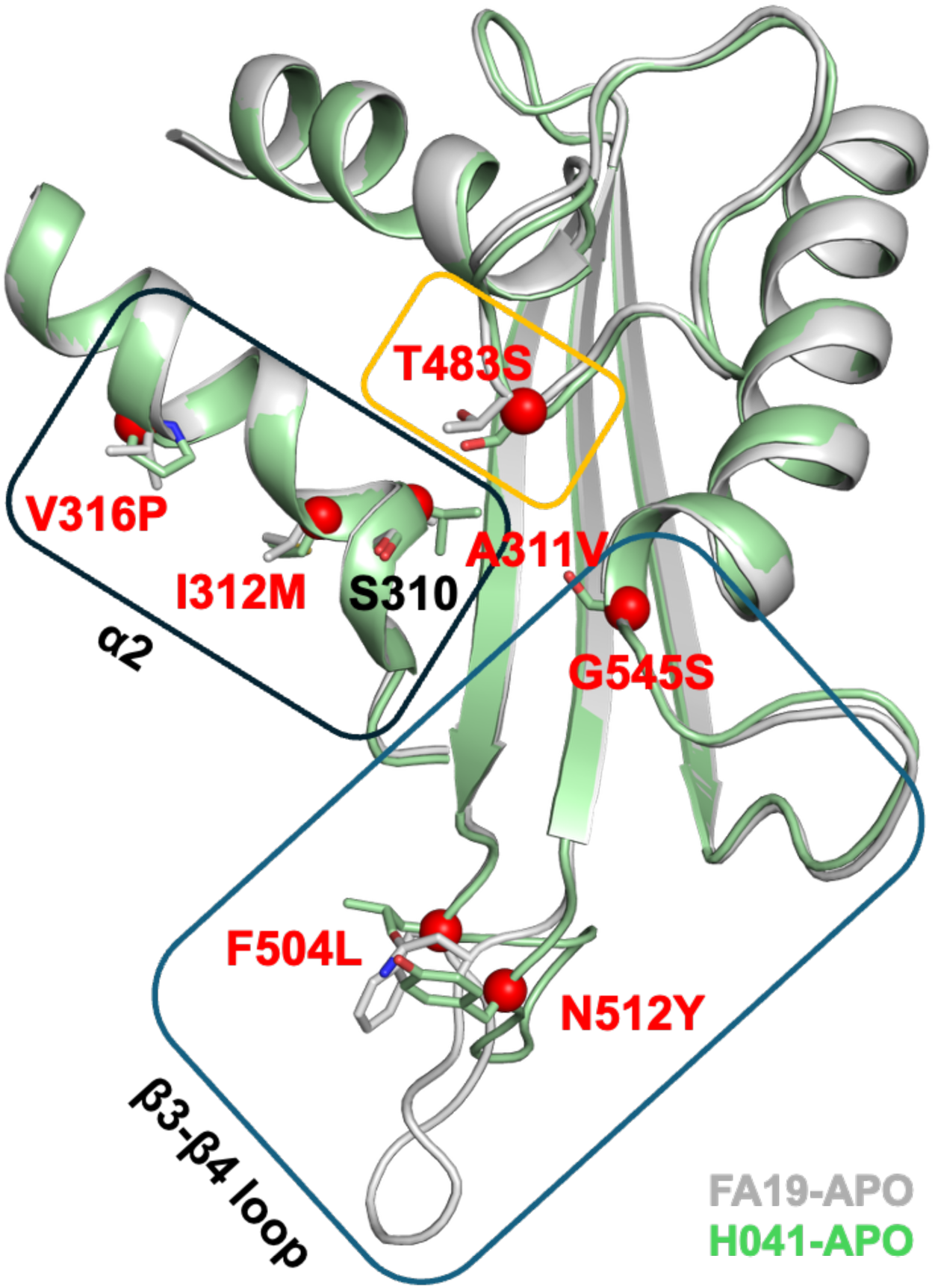
Mutations in PBP2 that confer ESC resistance to *N. gonorrhoeae*. A superimposition of two *apo* PBPs structures in the active site region is shown in ribbon format, where PBP2^FA19^ is colored grey and PBP2^H041^ is green. Boxes indicate the groups of mutations that were analyzed in this study. The side chains for resistance mutations are shown in stick form and colored according to structure. Red spheres indicate the alpha carbon position of these residues in PBP2^H041^. The Ser310 nucleophile is also shown. Note the different conformation of the 𝛽3-𝛽4 loop in the two structures.

## Results

In previous studies, we and others identified seven mutations in mosaic PBP2 variants that confer resistance to the ESCs ceftriaxone and cefixime [33, 35–38]. These mutations are clustered around the active site and can be grouped based on their location and/or effects on conformation: a) two 𝛽3-𝛽4 loop mutations (F504L/N512Y) and G545S, b) three mutations (A311V/I312M/V316P) located on the same 𝛼2 helix as the active site Ser310 nucleophile, and c) the T483S mutation at the top of the active site (Fig. 1). However, it is unclear whether these mutations act independently or not to decrease acylation with specific β-lactam antibiotics. To investigate potential dependencies, we introduced each of the groups of mutations into the *penA* gene from FA19 (*penA19*) or reverted them back to their *penA19* counterparts in the *penA* gene from H041 (*penA41*), transformed the mutant alleles into the antibiotic-susceptible strain, FA19, and determined the MICs of ceftriaxone, cefixime, and penicillin G.

### 𝛽3-𝛽4 loop mutations work in tandem with G545S to elevate MICs of ESCs

Introduction of either F504L/N512Y or G545S into *penA19* increases the MICs of ceftriaxone (MIC_CRO_) and cefixime (MIC_CFX_) by 1.5- to 3-fold, with G545S having a greater impact on the MICs than the 𝛽3-𝛽4 loop mutations (Fig. 2A). When all three mutations were introduced together, the MIC_CRO_ and MIC_CFX_ increased 3.5-fold and 6-fold, respectively, compared to *penA19*. Surprisingly, these mutations had no impact on the MIC of penicillin G (MIC_PEN_).

**Figure 2.**
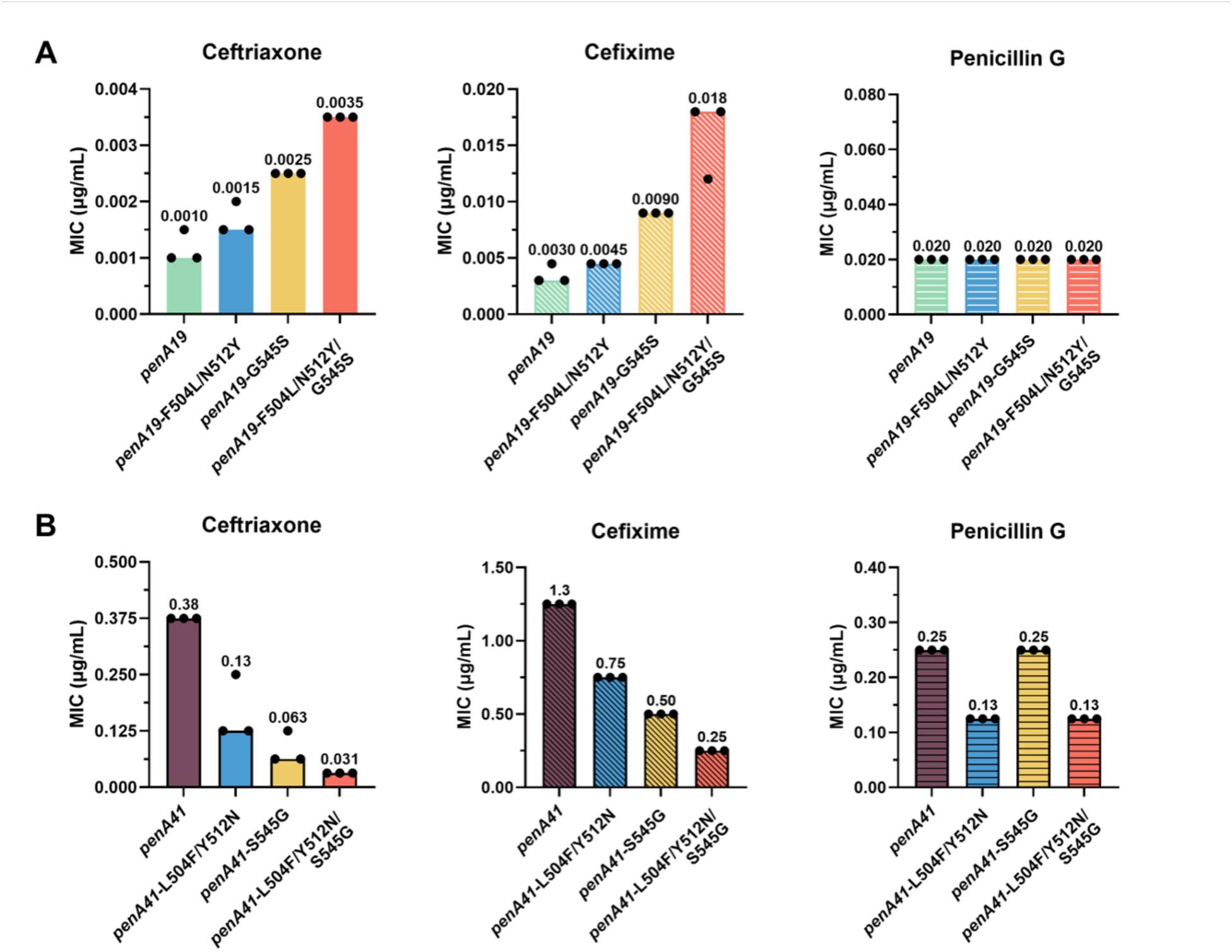
Effects of β3-β4 loop mutations and G545S on antibiotic MICs. MIC values of ceftriaxone, cefixime, and penicillin G in FA19 strains transformed with *penA19* containing the specified mutations from *penA41* (forward; A) or reversion of these mutations in *penA41* back to their counterparts in *penA19* (reversion; B). The bars represent the median MICs with individual replicates shown as points (n = 3).

Reverting these codons to their *penA19* equivalents in *penA41* showed a similar pattern, again with antibiotic-specific effects (Fig. 2B). Reverting Ser545 to Gly in *penA41* results in a larger decrease in the MIC_CRO_ and MIC_CFX_ than does reversion of the two 𝛽3-𝛽4 loop mutations. In contrast, reversion of Ser545 back to Gly had no effect on MIC_PEN_, whereas reverting the 𝛽3-𝛽4 loop mutations decreased MIC_PEN_ twofold. When compared to *penA41,* the MIC_CRO_ was decreased the most (12-fold) when all three mutations were reverted, followed by MIC_CFX_ (5-fold), and MIC_PEN_ (2-fold).

### Mutation and reversion of 𝛼2 helix amino acids markedly impact the MICs of ceftriaxone, cefixime, and penicillin G

Because of their proximity to one another, the three 𝛼2 mutations were mutated as a group to assess their contributions to ceftriaxone, cefixime, and penicillin G resistance. Mutation of the three codons to their *penA41* equivalents in *penA19* and transformation into FA19 increased the MIC_CRO_ and MIC_CFX_ by 5-and ∼7-fold, respectively, compared to FA19, but had little to no impact on the MIC_PEN_ (Fig. 3). In contrast, reverting these mutations to their wild-type residues in the *penA41* background had a marked impact on the MICs of all three antibiotics tested, with the MICs for ceftriaxone, cefixime, and penicillin G decreased by 6-fold, 15-fold, and 6-fold, respectively (Fig. 3). Notably, reversion of the three 𝛼2 mutations in *penA41* decreased the MIC_PEN_ to the same level as FA19.

**Figure 3.**
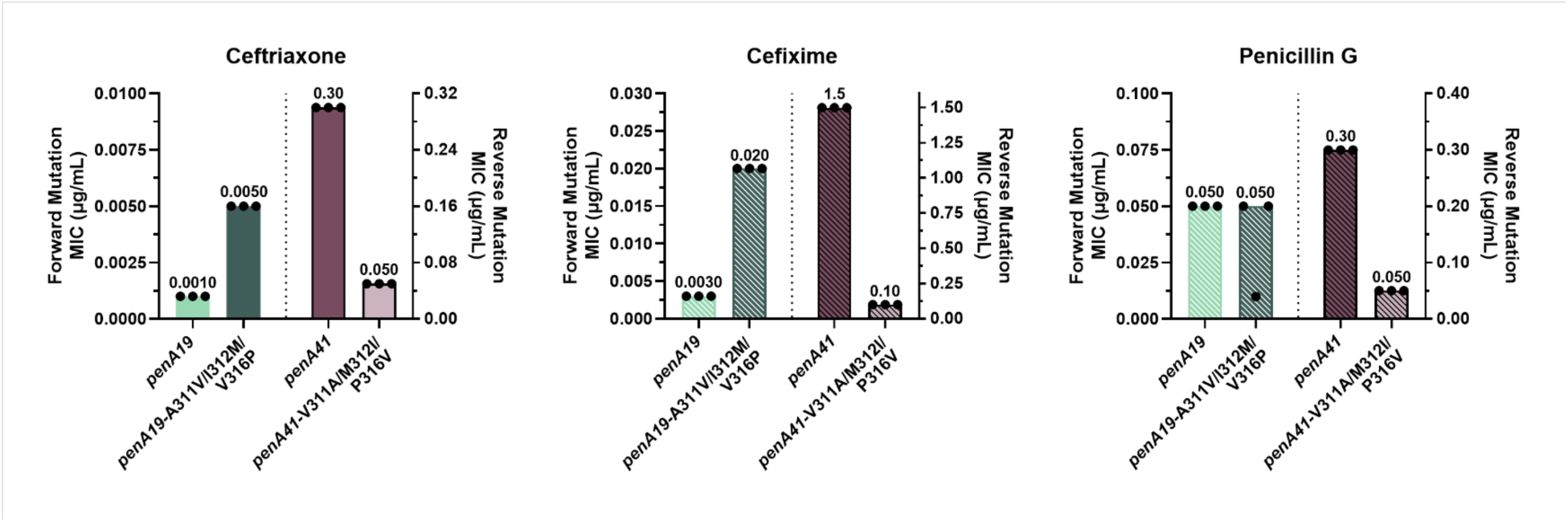
MIC values of ceftriaxone, cefixime, and penicillin G for FA19 strains transformed with *penA19* containing the 𝛼2 mutations from *penA41* (left side of each panel) or *penA41* containing reversions of amino acids 311, 312, and 316 to their counterparts in *penA19* (right side of each panel). Note that the left sides of each panel have a different Y axis than the right sides, due to the large differences in MICs of the antibiotics for the strains. The bars represent the median MICs with individual replicates shown as points (n = 3).

### T483S mutation

The T483S mutation sits at the top of the active site (Fig. 1) and plays an important role in resistance in H041 and other high-level ESC-resistant strains such as FC428 [37, 39]. In ESC-bound crystal structures of tPBP2^H041^ and tPBP2^FA19^, the Ser or Thr side chain at position 483 projects toward ceftriaxone, though does not form a direct contact with the antibiotic in either structure [32, 33]. We mutated Thr483 to Ser in *penA19* and attempted to transform FA19, but no colonies were obtained despite repeated attempts. In contrast, reversion of Ser483 to Thr in the *penA41* background decreased the MIC_CRO_ by 4-fold, along with MIC_CFX_ and MIC_PEN_ by 2- and 1.5-fold, respectively (Fig. 4).

**Figure 4.**
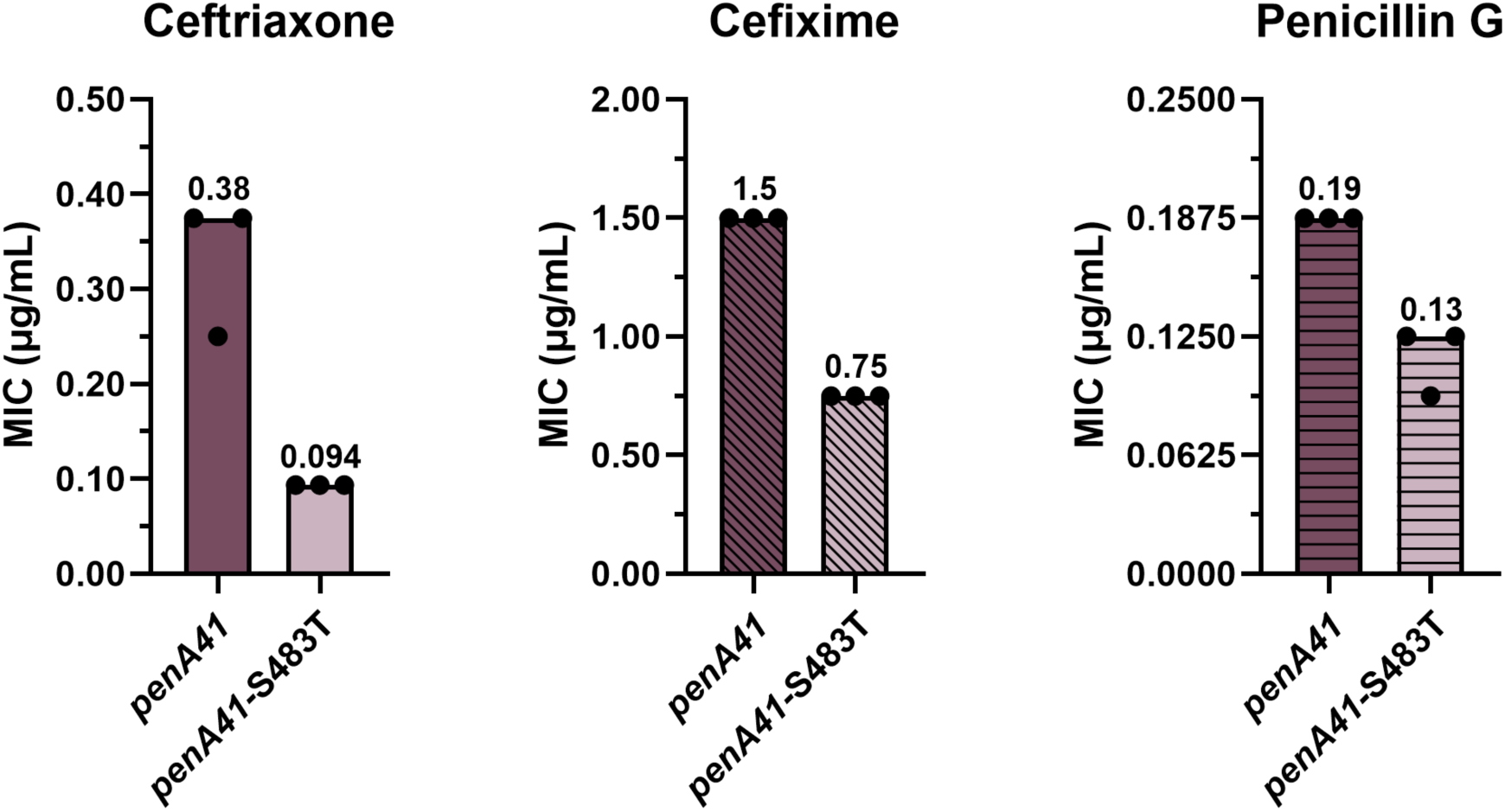
MIC values of ceftriaxone, cefixime, and penicillin G for FA19 strains transformed with *penA41* containing an S483T reversion. The bars represent the median MICs with individual replicates shown as points (n = 3).

### T483S disrupts transpeptidase function in the PBP2^FA19^ background without three epistatic mutations

To gain a measure of how much of the resistance to ESCs conferred by the full *penA41* allele depends on these seven identified resistance mutations, we initially introduced the two 𝛽3-𝛽4 loop mutations, G545S, and the three 𝛼2 mutations into *penA19* (referred to as *penA19*-6M), transformed the construct into FA19, and determined the MICs of the three β-lactam antibiotics. For the ESCs, the MIC_CFX_ reached 33% of that conferred by the *penA41* allele in FA19, whereas the MIC_CRO_ and MIC_PEN_ reached 25% and 16% of that conferred by *penA41*, respectively (Fig. 5). Unexpectedly, when we added the T483S mutation to *penA19*-6M (termed *penA19*-7M) and transformed it into FA19, only a few colonies were obtained on selection plates and all lacked the T483S mutation, suggesting that in the *penA19*-7M background, the T483S mutation disrupted the essential transpeptidase activity of PBP2 to nonviable levels. This conclusion also was consistent with the inability to select for strains harboring *penA19*-T483S described above.

**Figure 5.**
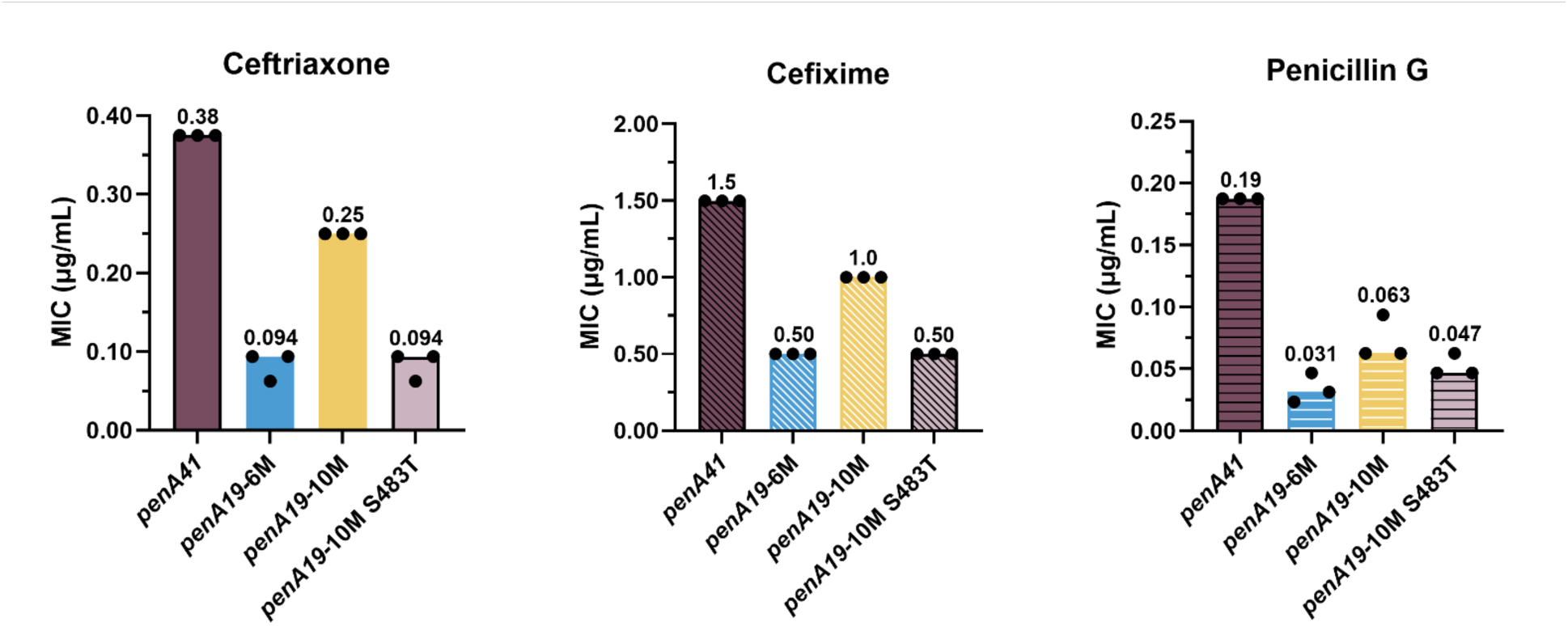
MIC values of ceftriaxone, cefixime, and penicillin G for FA19 strains transformed with *penA19* alleles containing multiple resistance mutations from *penA41*. The *penA19*-6M allele includes F504L, N512Y, G545S, A311V, I312M, and V316P resistance mutations. The *penA19*-10M allele contains all of the mutations in *penA19*-6M, plus T483S and the 3 epistatic mutations, A437V, L447V, and F462I. The bars represent the median MICs with individual replicates shown as points (n = 3).

We surmised that some of the other mutations in *penA41* were important in preserving transpeptidase activity in the presence of T483S. In previous studies on PBP2 from an ESC-reduced susceptibility *N. gonorrhoeae* strain (35/02) and ESC-resistant H041, we had divided the linear sequence into 6 modules to assess their contributions to resistance [36, 37] and a similar approach was used here to identify these additional mutations. When the various modules (mods) from *penA41* were inserted into *penA19*-7M, only mod3^H041^ produced transformants. Sequencing of isolates confirmed that all seven of the resistance mutations, including T483S, were intact. Mod3 encompasses residues 432-489 and contains T483S as well as 12 additional mutations. By trial and error (see Supplemental Results, Fig. S1, and Table S1 for a detailed description), we winnowed down the twelve mod3^H041^ mutations to three-A437V, L447V, and F462I-that together supported growth in *penA19*-7M (this construct is referred to as *penA19*-10M). Mapping these onto the structure of PBP2 shows they are relatively distant from the active site region (Fig. 6). The MICs of ceftriaxone and cefixime for FA19 *penA19*-10M reached ∼67% of that of FA19 *penA41*, with a more moderate impact on the MIC_PEN_ (Fig. 5). Importantly, when Ser483 in *penA19*-10M was reverted to Thr and transformed into FA19 (*penA19*-10M S483T), the MICs of ceftriaxone and cefixime decreased to the same level as that conferred by *penA19*-6M (Fig. 5). Taken together, these data reveal that the A437V, L447V, and F462I mutations have an epistatic effect on the T483S mutation, meaning that they do not contribute to resistance to ESCs but are required for the *penA19*-7M mutant to be viable.

**Figure 6.**
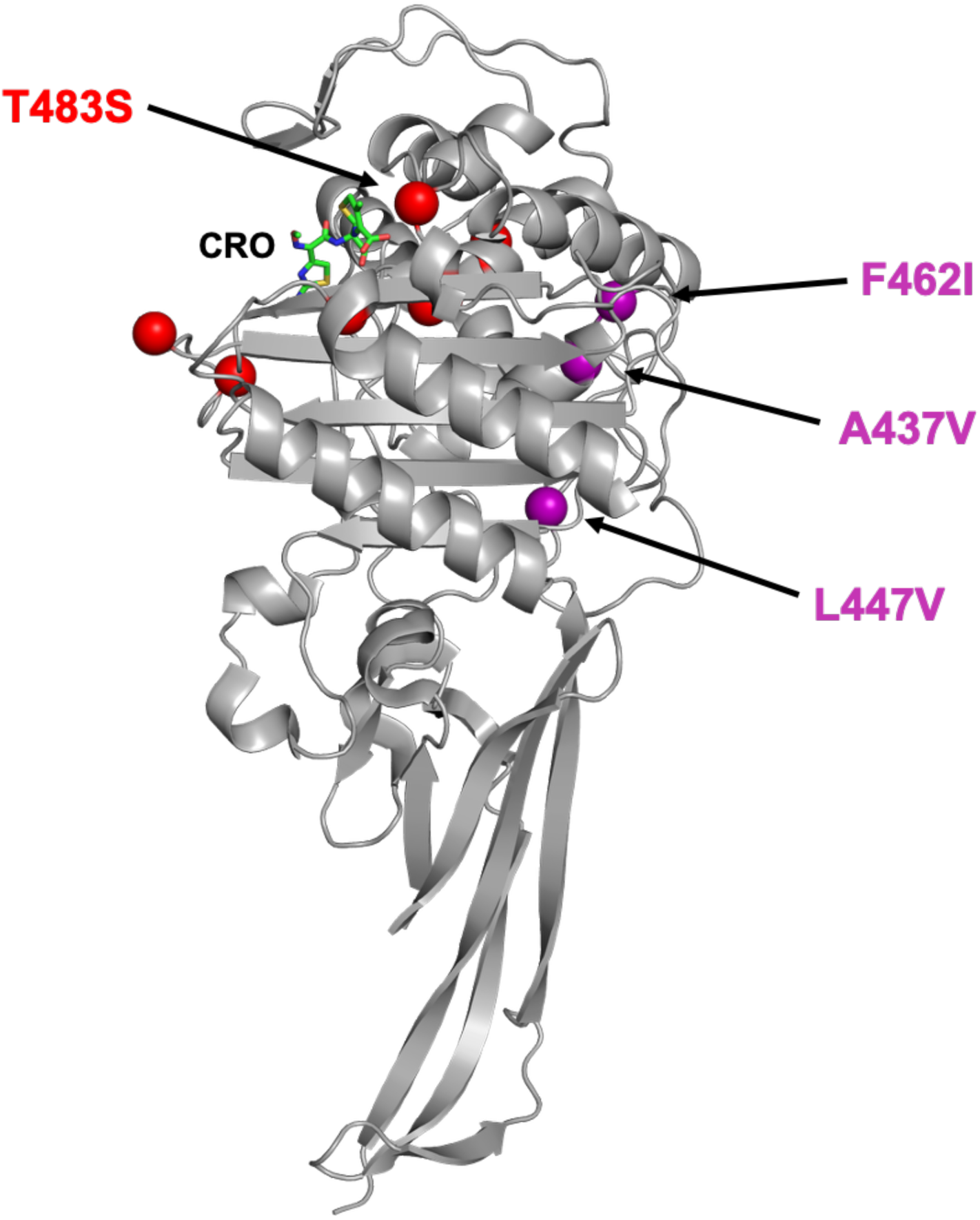
Location of the three epistatic mutations in PBP2. The epistatic mutations are marked by purple spheres and the seven resistance mutations around the active site are shown by red spheres. CRO indicates the location of ceftriaxone (green bonds) bound in the active site.

### Acylation kinetics of PBP2 variants

To assess the *in vitro* effects of T483S on antibiotic binding to PBP2 in the presence or absence of the three epistatic mutations, we determined the *k_2_*/K_s_ values of PBP2-6M, PBP2-7M, and PBP2-10M using purified proteins and compared them to previously reported values for PBP2^FA19^ and PBP2^H041^. As shown in Table 1, the three antibiotics have extremely high *k_2_*/K_s_ values for PBP2^FA19^, whereas these values are markedly lower for PBP2^H041^, consistent with their respective MICs for FA19 and H041. The *k_2_*/K_s_ values of penicillin G, ceftriaxone, and cefixime are decreased by 38-fold, 620-fold, and 84-fold, respectively, for PBP2-6M relative to those for PBP2^FA19^, and adding the T483S mutation (PBP2-7M) lowers them by another 7- to 13-fold. The *k_2_*/K_s_ values of the three antibiotics for PBP2-10M are very similar to those for PBP2-7M, consistent with the lack of an effect of epistatic mutations on antibiotic binding.

**TABLE 1.** *k_2_*/K_s_ constants of Penicillin G, Ceftriaxone, and Cefixime for PBP2-6M, PBP2-7M, and PBP2-10M. *k_2_*/K_s_ values were derived using [^14^C]benzyl penicillin G as described in [24]. ^a^Data from [37].

| | $k_2/K_s$ ($\text{M}^{-1}\text{s}^{-1}$) | | |
| --- | --- | --- | --- |
| **PBP2 Protein** | **Penicillin G** | **Ceftriaxone** | **Cefixime** |
| <sup>a</sup>PBP2<sup>FA19</sup> | 75,700 ± 2,300 | 1,710,000 ± 90,000 | 1,480,000 ± 22,000 |
| PBP2-6M | 2,000 ± 43 | 2,745 ± 140 | 17,550 ± 644 |
| PBP2-7M | 285 ± 15 | 305 ± 7 | 1,343 ± 54 |
| PBP2-10M | 352 ± 12 | 483 ± 24 | 2,103 ± 45 |
| <sup>a</sup>PBP2<sup>H041</sup> | 741 ± 28 | 135 ± 21 | 55 ± 14 |

### The three epistatic mutations are insufficient to maintain transpeptidase activity when incorporated into penA19-T483S

The data detailed above suggested that one potential reason we could not select colonies following transformation of *penA19*-T483S into FA19 was because the inclusion of T483S disabled the essential transpeptidase activity of PBP2. We tested this idea by introducing the three epistatic mutations into *penA19*-T483S and transforming the construct into FA19. We obtained a small number of transformants (eight total in two experiments), but while we were able to incorporate the T483S mutation into FA19, surprisingly all the transformants also contained a spontaneous R502H mutation. Arg502 is 100% conserved in all sequenced *penA* alleles in *N. gonorrhoeae* [40], is located next to the highly conserved KTG sequence motif (residues 497-499) on the β3 strand [22], and contacts both ceftriaxone and cefixime in the acylated structures of tPBP2^H041^ [32, 33, 38]. Thus, even with the three epistatic mutations present, an additional non-native mutation (R502H) is necessary for the T483S mutation to maintain sufficient transpeptidase activity for growth in the *penA19* background.

### Epistatic mutations correlate with T483S in Neisseria species

Given the severe effects of the T483S mutation on PBP2 transpeptidase activity, we examined its prevalence in *N. gonorrhoeae*, *N. meningitidis*, and commensal *Neisseria* species in the PubMLST database [40]. In *N. gonorrhoeae*, 0.7% (216 out of 32,719) of genomes within the database contained the T483S variation, likely representing strains with mosaic *penA* alleles. *Neisseria meningitidis*, the etiological agent of meningococcal disease, has less ESC-reduced susceptibility than *N. gonorrhoeae*, but recent studies have documented the emergence of such strains [41, 42]. In the PubMLST database, 0.07% (34 out of 45,753) of meningococcal genomes contained the T483S mutation. Because mosaic *penA* alleles are known to arise through recombination events with commensal *Neisseria* species [29], we also determined the prevalence of Thr483 within 40 commensal *Neisseria* species in the PubMLST database. Across those 40 strains, 1.4% (25 out of 1,802) of genomes contained the T483S mutation. We also examined the conservation of the three epistatic mutations in *penA* alleles from all *Neisseria* species that contained a T483S mutation. Of these, the epistatic mutations A437V, L447V, and F462I were observed in 98% of sequences, consistent with the requirement of the epistatic mutations to preserve essential transpeptidase activity in variants harboring a T483S mutation.

### Structural comparison of minimal mutant PBP2 variants

To assess the molecular impact of these mutations on PBP2, we determined the crystal structures of the tPBP2-6M, -7M, and - 10M variants. All crystals grew in the same P2_1_ crystal system as tPBP2^FA19^ [32] with two molecules in the asymmetric unit, except that tPBP2-10M crystallized with slightly altered cell dimensions (Table S2). All three structures superimpose closely with each other and with the structures of tPBP2^FA19^ and tPBP2^H041^, with RMS deviations in the range 0.17 to 0.47 Å for all atoms (Fig. 7A). Only the 𝛽3-𝛽4 loop differs structurally, occupying an extended conformation in the tPBP2-6M, -7M, and -10M variants similar to that in tPBP2^FA19^, whereas it occupies a so-called “outbent” conformation in tPBP2^H041^ (Fig. 7B). This is interesting because the 6M, 7M and 10M variants all contain the F504L and N512Y mutations, showing that in *apo* structures at least, these mutations do not affect the conformation of the 𝛽3-𝛽4 loop.

**Figure 7.**
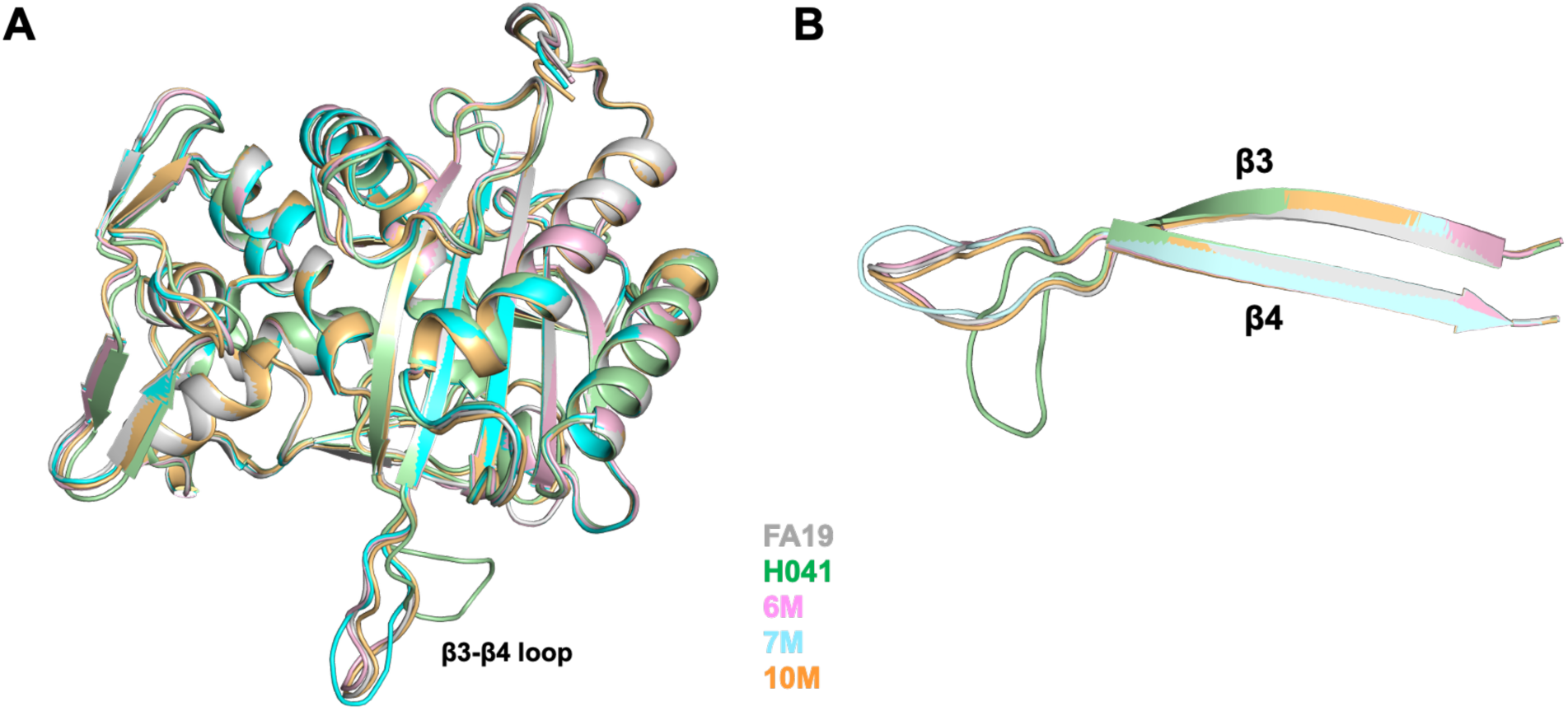
Epistatic mutations in PBP2 do not elicit major changes in structure. **A.** Superimposition of the transpeptidase domains of PBP2 structures. **B.** Close-up of the 𝛽3-𝛽4 loop, showing its extended conformation in tPBP2^FA19^, tPBP2-6M, tPBP2-7M and tPBP2-10M. This contrasts with its “outbent” conformation in PBP2^H041^.

Examination of the structures around the sites of epistatic mutations reveals some interesting findings. Firstly, all three mutations are hydrophobic to hydrophobic substitutions, where two side chains have become smaller (L447V and F462I) and one larger (A437V), and all three are situated within hydrophobic core regions of the protein. Secondly, superimpositions of the variant structures with that of tPBP2^FA19^ and tPBP2^H041^ show that the hydrophobic core regions around the F462I and A437V mutations are essentially unchanged, whereas in tPBP2-10M, the structure is altered around the L447V mutation (Fig. S2). Specifically, residues on the 𝛼6-𝛽2e loop are altered in position, as is the side chain of Phe499. This points to the L447V mutation as having the biggest effect structurally. Finally, a closer examination shows that the dihedral angles for the 446-447 peptide bond have switched from being a Ramachandran outlier in the tPBP2-6M and tPBP2-7M structures to the allowed region in tPBP2-10M (Fig. 8). This occurs in both molecules of the asymmetric unit. Interestingly, this bond is a Ramachandran outlier in tPBP2^FA19^ but occupies the allowed region in tPBP2^H041^. This indicates that the L447V mutation has relieved strain present in structures that lack this mutation. Taken together, these data demonstrate that the epistatic mutations have an identifiable effect on protein structure that allows T483S to retain sufficient transpeptidase activity to remain viable.

**Figure 8.**
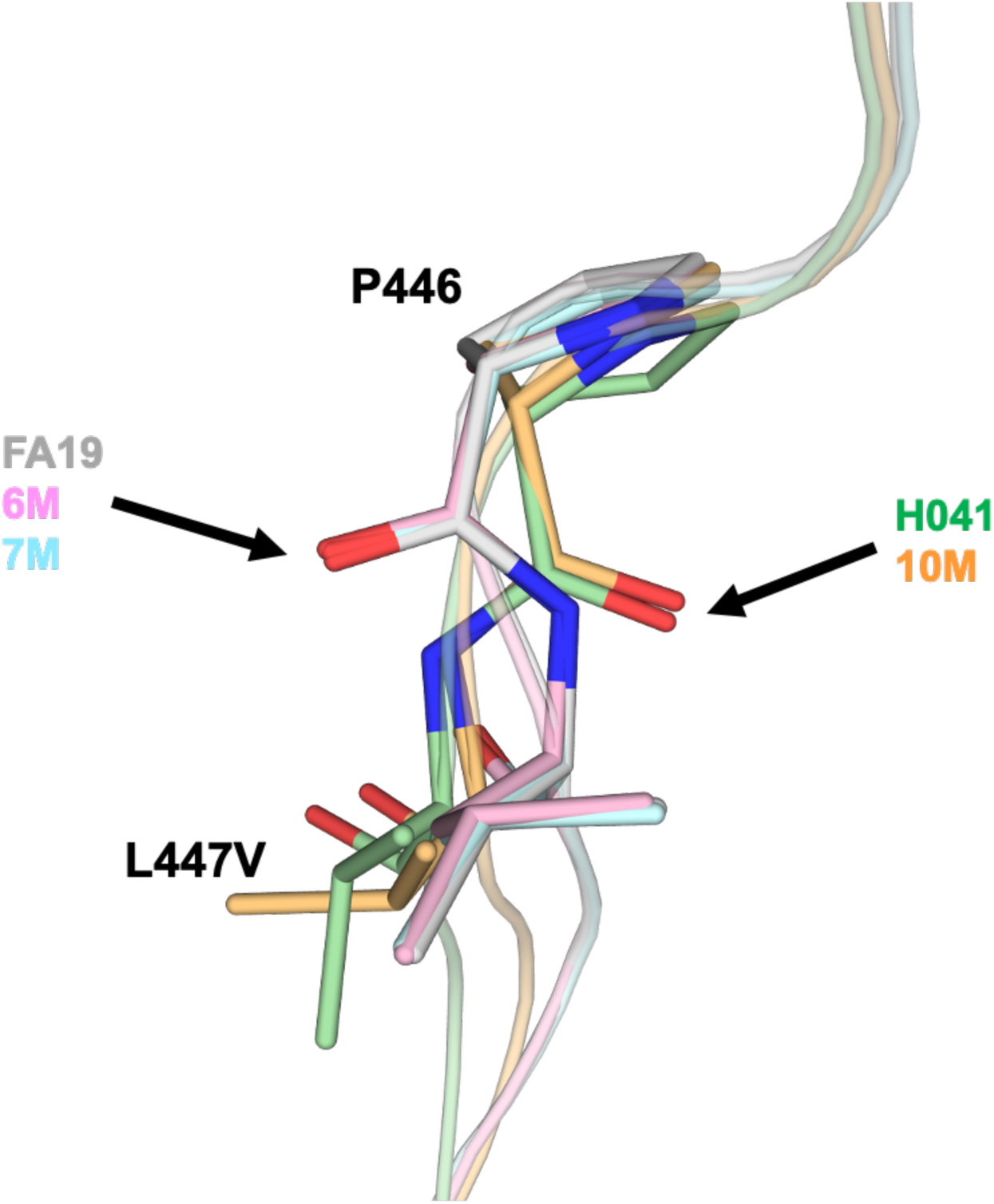
Stereochemistry of the peptide bond between 446 and 447 is altered by epistatic mutations. As marked by arrows, the dihedral angles for 446-447 occupy a disallowed region in the crystal structures of tPBP2^FA19^, tPBP2-6M and tPBP2-7M, but these occupy the favorable region in PBP2-10M and PBP2^H041^. Structures are colored the same as Fig. 7.

### Growth deficits by key resistance mutations are overcome by fitness-promoting mutations

Our data with the T483S mutation suggest that resistance mutations in PBP2 can negatively impact the essential transpeptidase function of the enzyme and thereby decrease the fitness of strains. To examine how the different resistance mutations impact strain growth (a proxy for biological fitness), we carried out quantitative growth curves of FA19 harboring key *penA* variants (Fig. 9). Surprisingly, across all eight hours of growth, there were no significant differences in OD_600_ readings between FA19 *penA19* and FA19 *penA41* (Fig. 9B) at any time (Table S3). In contrast, FA19 *penA19-*10M had significantly impaired growth compared to FA19 *penA19* at 4 hours and both FA19 *penA41* and FA19 *penA19* at 8 hours (Fig. 9B-C), suggesting that while the three epistatic mutations repair the transpeptidase function damaged by T483S, the enzyme is still impaired. The T483S mutation imparts marked negative effects on fitness, as FA19 *penA19*-10M S483T (the same as the 6M construct plus three epistatic mutations) showed significantly higher growth at the 8-hour timepoint compared the FA19 *penA19*-10M (Fig. 9C) and FA19 *penA19*-6M (Fig. 9D). There was no significant difference between *penA19* 10M S483T and *penA41* (Fig. 9D).

**Figure 9.**
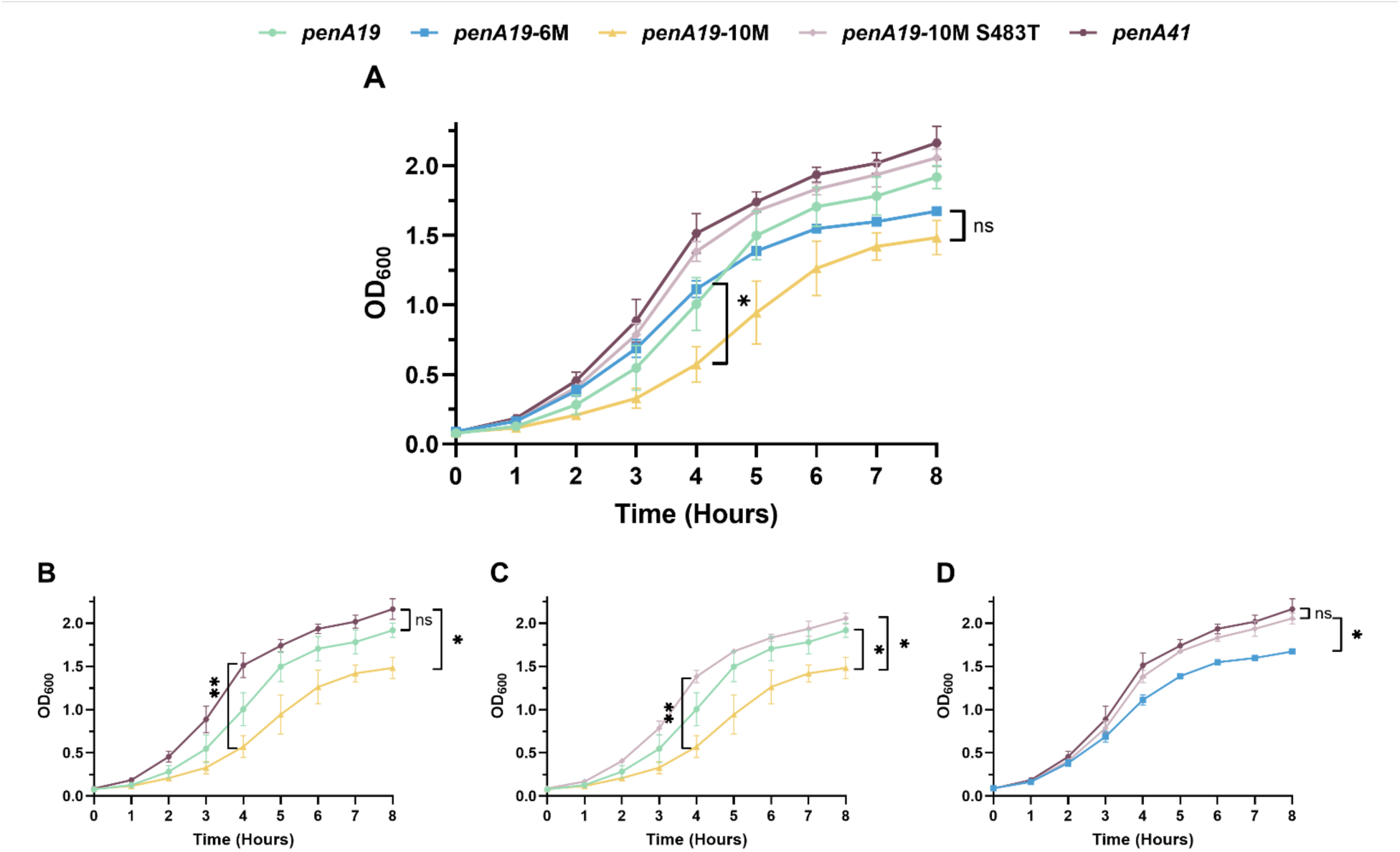
Quantitative growth curves of individually cultured FA19 strains with the indicated *penA* alleles. Legend for lines and symbols shown above graphs. **A.** Overlay of growth curves from all five strains tested. n = 3, with error bars showing standard deviation. At the 8-hour time point, there is no significant difference in OD_600_ between *penA19* 6M and *penA19* 10M (p > 0.05). B-D. Overlay of growth curve from the indicated strains. B. At the 8-hour time point, there is a significant (p = 0.011) difference in OD between FA19 *penA41* and FA19 *penA19-*10M strains. At all time points tested, there is no significant difference between *penA19* and *penA41* (p > 0.05). **C.** At the 8-hour time point, there is a significant difference in OD when comparing FA19 *penA19-*10M to both FA19 *penA19* (p = 0.042) and FA19 *penA19-*10M S483T (p = 0.022). **D.** At the 8-hour time point, there is a significant (p = 0.022) difference in OD_600_ between FA19 *penA19-*6M and FA19 *penA19-*10M S483T strains. Statistical significance was calculated using repeated-measures 2-way ANOVA with Geisser-Greenhouse correction and Tukey’s multiple comparisons.

## Discussion

In this study, we set out to understand whether the resistance mutations in the *penA41* allele from the ESC-resistant strain H041 act independently or depend on one another for their capacity to confer resistance to three β-lactam antibiotics that have been used historically to treat gonorrhea infections. By clustering the seven known resistance mutations into three groups and assessing both forward mutagenesis (incorporating mutations into the antibiotic-susceptible *penA19* allele) and reverse mutagenesis (reverting mutations to wild-type in the antibiotic-resistant *penA41* allele), we determined that these mutations work together to increase resistance, but the results differed depending on which antibiotic was being tested. Overall, we observed that the MICs of the ESCs ceftriaxone and cefixime were more strongly affected by the different groups of mutations than that of penicillin G, suggesting that the mutations we examined are more important for ESC resistance than penicillin G resistance. We also uncovered a unique dependence of one of the mutations, T483S, on three epistatic mutations in a region on the opposite side of the molecule that have no impact on resistance but are required for maintaining sufficient transpeptidase activity to support growth. The crystal structures of tPBP2-6M, -7M, and -10M revealed that the unfavorable backbone dihedral angles at residues 446-447 are relieved by the three epistatic mutations in tPBP2-10M, suggesting that this conformational change is important in supporting transpeptidase activity of PBP2 variants containing the T483S mutation. Lastly, quantitative growth curves show that despite the presence of the epistatic mutations, the *penA19*-10M allele still carries a strong fitness cost, because reversion of T483S back to wild-type increases the growth of the strain to the same levels as FA19 and FA19 *penA41*. Taken together, these data suggest a complex interplay between mutations that increase resistance with other mutations that play a supporting role in maintaining essential transpeptidase activity.

Our results revealed two interesting observations. First, there was a marked difference between the impact of the mutations on the MICs of the ESCs relative to the MIC_PEN_. For example, incorporating the 𝛽3-𝛽4 loop mutations plus G545S or the three 𝛼2 mutations into *penA19* had no effect on MIC_PEN_, whereas the MICs of CRO and CFX were increased 3.5- to 7-fold. The lack of an effect of the seven mutations examined here on MIC_PEN_ suggests that they are ESC-specific, and that other mutations in *penA41* are involved in increasing the MIC_PEN_. This is perhaps not unexpected given the structural differences in these antibiotics and the fact that the *penA41* allele evolved to confer resistance to ESCs. Interestingly, whereas the 𝛼2 mutations had no effect on the activity of penicillin in the forward direction, reverting those mutations in *penA41* decreased the MIC_PEN_ by 6-fold. These data suggest that the 𝛼2 mutations may coordinate with the penicillin-specific mutations in *penA41* but by themselves do not affect the activity of penicillin G against PBP2. Secondly, although there was some variability, reverting mutations to wild-type in *penA41* had larger fold differences in the MICs for ESCs compared to those when introducing them into *penA19*. This hints at synergism between mutations in the context of *penA41* that do not occur in the *penA19* background.

The capacity of a 𝛽-lactam antibiotic to inhibit a PBP is defined by the second-order rate of acylation, *k_2_*/K_s_, in which *k_2_* is the rate of acylation and K_s_ is the non-covalent binding affinity (equivalent to K_d_; [43]). Thus, resistance mutations can decrease the acylation rate constant *k_2_*, decrease the non-covalent binding affinity (i.e. increase K_s_), or both. We have previously shown by isothermal titration calorimetry that ceftriaxone binds to tPBP2^FA19^-S310A, an acylation-incompetent mutant, with a K_s_ value in the low-micromolar range, but not to tPBP2^H041^-S310A [33]. We also showed that introducing the 𝛽3-𝛽4 loop mutations individually into tPBP2^FA19^-S310A increased the K_s_ of ceftriaxone 2- to 4-fold, G545S increased K_s_ by 10-fold, and introducing all three together increased the K_s_ >100-fold [38]. Thus, a good portion of the decrease in the MIC_CRO_ in *penA19*-F504L/N512Y/G545S is likely due to the increase in K_s_, although these mutations (particularly G545S) may also decrease *k_2_* for reasons described for the 𝛼2 mutations below.

In contrast to the 𝛽3-𝛽4 loop mutations and G545S, the 𝛼2 mutations are unlikely to affect binding, since the position of the 𝛼2 helix is identical in *apo* and acylated PBP2 crystal structures [32, 33]. Instead, their effects on the MICs of ESCs in the forward direction are likely due to decreasing *k_2_*. We suspect that the 𝛼2 helix moves towards the 𝛽3 strand during the development of the transition state and concomitant formation of the oxyanion hole (comprised of the main chain amides of Ser310 and Thr500) and then relaxes back to its original location following acylation. It therefore seems likely that these three mutations hinder the transient movement of the 𝛼2 helix during acylation, thereby lowering *k_2_* and decreasing acylation.

Incorporating all seven resistance mutations and the three epistatic mutations into *penA19* increased the MICs of the ESCs to 67% of those conferred by *penA41*. Identifying the remaining resistance mutations in *penA41* will be difficult for multiple reasons: a) in the absence of the mosaic background, such mutations might lower transpeptidase activity to the level that no longer supports growth; b) the changes in MIC with additional mutations may be incremental, making it difficult to select; and/or c) it may require the presence of multiple mutations that work together to increase the MIC. Nonetheless, showing the high level of resistance conferred by ten mutations (of which three do not increase resistance alone but repair the transpeptidase activity) from a total of 60 mutations in *penA41* relative to *penA19* is an important achievement and provides a framework for understanding the complexities of resistance conferred by mutations in PBP2.

It is important to note that while most of the mutations identified here are present in other mosaic alleles, there are some important differences. For example, FC428 is an internationally spreading ceftriaxone-resistant clone that has a lower MIC_CRO_ than H041 (0.5 µg/mL vs 2-4 µg/mL for H041) [39]. FC428 harbors the *penA60* mosaic allele that contains 48 mutations compared to *penA19*; notably, six of the seven mutations investigated here are found in *penA60*, with the seventh (V316P) mutated to a different amino acid (V316T). Additionally, F89 (MIC_CRO_=1-2 µg/ml) lacks the T483S and V316P/T mutations but has an A501P mutation just downstream of the KTG motif [44]. It seems likely that the mosaic background is more conducive to acquiring new mutations than *penA19* or similar alleles, and single mutations may arise spontaneously in the presence of selective pressure that further increase resistance.

The T483S mutation is an enigma. In *penA19*-6M with the three epistatic mutations, it increases the MIC_CRO_ by ∼2.5-fold, and when reverted in *penA41*, decreases the MIC_CRO_ by 4-fold. Yet, for T483S to be tolerated by the cell in the *penA19*-6M background, it requires the three epistatic residues, presumably because they restore the transpeptidase activity of PBP2 that is otherwise lost when the T483S mutation is present. This is even more remarkable given that Thr to Ser is a highly conservative mutation, amounting to only loss of a methyl on the side chain. In antibiotic-bound structures of tPBP2^FA19^, Thr483 is close to the active site, where its side chain occupies the space between Thr498, the dihydrothiazine ring of ESCs, and Ser362 of the SxN motif (Fig. 10), and it forms a hydrogen bond with Thr498. As noted earlier, Thr498 is important for catalysis because its side chain rotates to engage the carboxylate of bound ceftriaxone/cefixime and is also believed to trigger the inward movement of the 𝛽3-𝛽4 loop [32]. Interestingly, the same hydrogen bond can form when 483 is a Ser (*i.e.* in PBP2^H041^), provided the Ser498 side chain has undergone similar rotation and the 𝛽3-𝛽4 loop occupies the inward conformation, *i.e.* in the crystal structures of PBP2^H041^ in complex with cefoperazone or piperacillin [45] (Fig. 10). Hence, these data point to the absence of the methyl group in Ser483 as being key for resistance rather than the hydroxyl. As one potential mechanism for how it might contribute to resistance, increased flexibility of the side chain imparted by loss of the methyl makes formation of the hydrogen bond with Thr498 less favorable with Ser compared to Thr. In turn, this makes it more difficult to form the productive state for acylation by cephalosporins.

**Figure 10.**
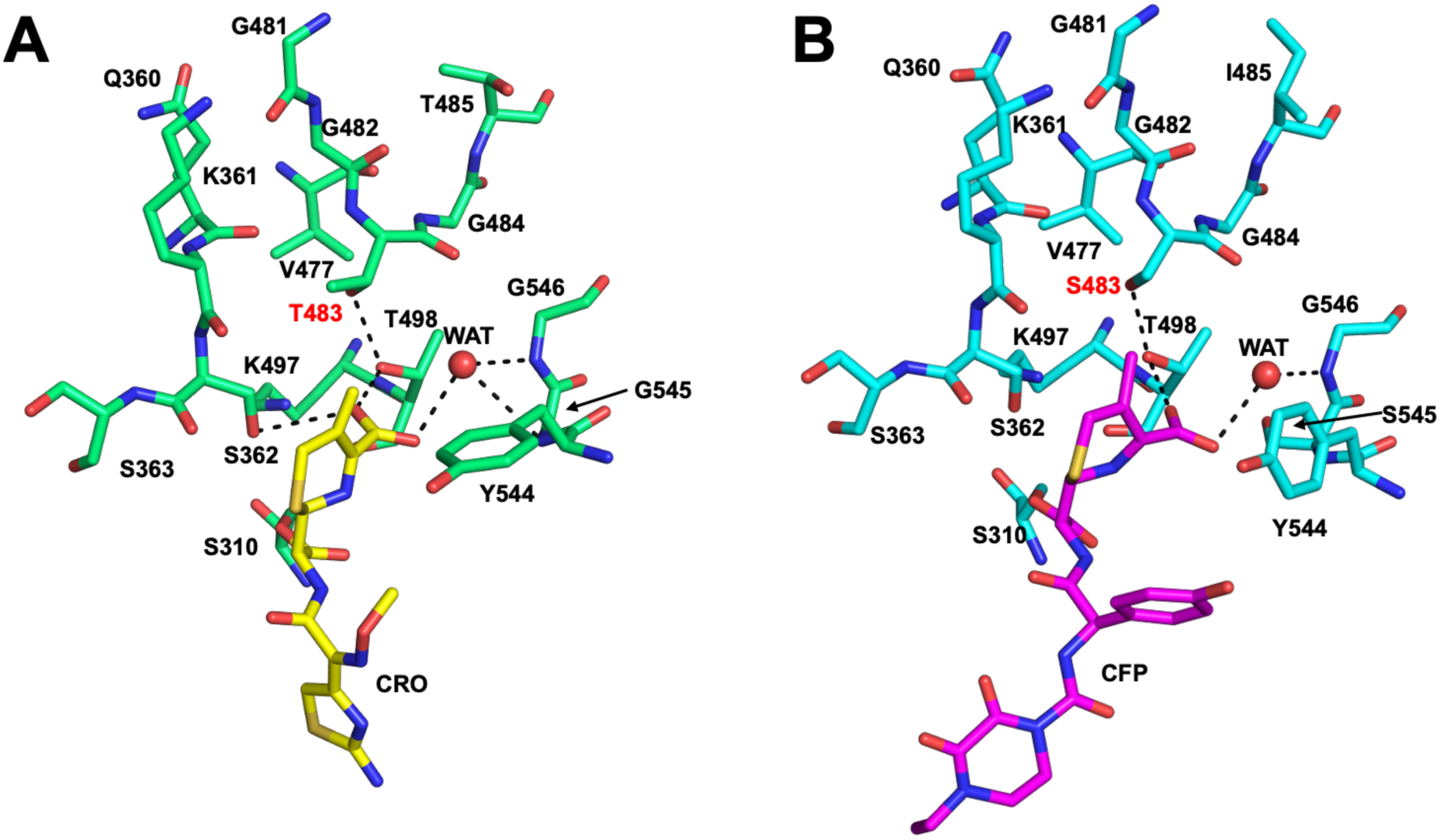
Thr483 and Ser483 have similar packing environments in crystal structures of PBP2 and the hydrogen bond with Thr498 is preserved. **A.** Environment around Thr483 in the crystal structure of PBP2^FA19^ (green bonds) bound by ceftriaxone (CRO, yellow bonds) [32]. **B.** Environment around Ser483 in the crystal structure of PBP2^H041^ (cyan bonds) bound by cefoperazone (CFP, purple bonds) [45]. For both panels, potential hydrogen bonds are indicated by dashed lines and waters are indicated by red spheres.

While it is possible to hypothesize how the T483S mutation can lower the rate of acylation, it is more challenging to understand how the three epistatic residues maintain transpeptidase function that would otherwise be compromised by a Ser at position 483. These mutations are located 22-28 Å away and therefore are unlikely to act through any direct contact. This suggests that they exert their effects allosterically via communication through the protein framework. It is interesting that all three residues are within hydrophobic cores, where alteration of side chains could perturb the surrounding environment. It is also notable that two of the mutations, A437V and F462I, pack within a hydrophobic core shared by α2, the helix that contains the Ser310 nucleophile at its N-terminal end. Presumably, any structural differences elicited by these two mutations are too small in magnitude to be detected by X-ray crystallography, because all structures overlap closely around these residues (Fig. S2). The situation is clearer for the L447V mutation, because the dihedral angles of the 446-447 peptide bond have switched from being in the unfavorable region of the Ramachandran plot in the structures of tPBP2^FA19^, tPBP2-6M and tPBP2-7M to being allowed in tPBP2-10M and the same as tPBP2^H041^ (Fig. 8). Consequently, there are some structural alterations around the L447V mutation, including shift of the 𝛼6-𝛽2e loop and of the Phe449 side chain (Fig. S2C). How these changes might affect Ser483 is unclear, however, since the position of this residue is similar in all structures (Fig. S3).

Quantitative growth curves in isogenic strains (with only the *penA* allele being altered) serve as a proxy for overall strain fitness and transpeptidase activity of PBP2 [46]. Surprisingly, we observed no significant difference in growth in GCB medium between FA19 and FA19 *penA41* across all timepoints tested, despite having shown previously that the *penA41* allele conferred a significant growth defect to FA19 [47]. One possibility for this difference is that the current experiments utilized a different lot of proteose peptone; indeed, the optical density at 600 nm for FA19 after 8 hours of growth reached >2, whereas it reached ∼1.0 in previous experiments. This difference may suggest that certain nutrients in different lots of proteose peptone #3 could alter growth of strains with mosaic alleles, although further experimentation to test this is warranted. Interestingly, [46] showed there was no difference in *in vitro* growth rates in GCB of isogenic strains harboring *penA10*, which is found in strains with reduced ESC susceptibility, or *penA60*, which is found in circulating strains such as FC428. However, the isogenic strain with *penA60* grew better than the strain with *penA10* when either palmitic acid or lithocholic acid was added to the growth medium. These data suggest a complex relationship between the *penA* allele and the composition of the growth medium that is not fully understood.

## Conclusion

In summary, the data presented here represent the first and only comprehensive analysis of resistance mutations and their role in essential transpeptidation in *N. gonorrhoeae* PBP2 and indeed in any highly remodeled PBP from other bacteria. The results indicate a complex and delicate balance between function and antibiotic resistance during remodeling of an essential enzyme. Because of the structural mimicry between β-lactam antibiotics and the acyl-D-Ala-D-Ala, the bacteria must subtly alter residues that can discriminate between antibiotic and substrate. These changes very often come with a cost in terms of enzymatic function and thereby exert a fitness cost, which is very evident from our results described here. It is likely that PBP-mediated ESC resistance at the level seen with *penA41* (∼400-fold increase compared to FA19) is unique to naturally competent organisms such as *N. gonorrhoeae*, as incorporating and maintaining multiple iterative resistance mutations in a wild-type background over time would be very difficult. This is also supported by the historical appearance of new mosaic *penA* alleles that confer increasing levels of resistance; once the mosaic background was present, the selective pressure exerted by ESCs drove additional mutations.

## Experimental Procedures

### Materials

Ceftriaxone sodium salt hemiheptahydrate and cefixime trihydrate were purchased from Thermo Fisher. Penicillin G sodium salt was purchased from Millipore Sigma. Other reagents such as culture media, buffers, and salts were purchased from commercial vendors.

### N. gonorrhoeae strains and culturing

FA19 is an antibiotic susceptible strain of *N. gonorrhoeae* that was used as the recipient strain for our transformation experiments [48]. H041 is a highly ESC-and penicillin-resistant strain that was isolated in Japan in 2009 [30, 31]. Strains were grown on Gonococcal Base Medium+ (GCB+) agar plates, which were made by supplementing GC Medium Base (Difco^TM^, BD) with Kellogg’s supplements I (4 mg/mL glucose, 0.1 mg/mL L-glutamate, and 0.2 µg/mL cocarboxylase, final concentrations) and II (5 µg/mL Fe(NO_3_)_3_) [49], and an additional 0.125% (w/v) agar (Bacto^TM^, BD). For *N. gonorrhoeae* growth in liquid media, GCB broth (15 g/L Proteose Peptone #3 (Bacto^TM^, BD), 4 g/L K_2_HPO_4_ (Fisher), 1 g/L potassium phosphate, monobasic (KH_2_PO_4_, Fisher), 5 g/L NaCl (Fisher), and 1 g/L starch from potato, soluble (Sigma)) was supplemented with 10 mM MgCl_2_, 20 mM sodium bicarbonate, and Kellogg’s supplements I and II to make liquid GCB+. For quantitative growth curves, liquid GCB+ without sodium bicarbonate was used and cultures were grown in a 37 °C incubator with 5% CO_2_ in a 50 mL tissue culture flask fitted with a gas-permeable cap.

### Transformation of FA19

Piliated (P+) *N. gonorrhoeae* is naturally competent, and transformation can be achieved simply by mixing DNA and P+ cells and plating on selective medium after incubating for 5 hrs as previously described [37]. All transformants were selected on GCB agar plates containing cefixime as the selection antibiotic. The concentrations differed depending on the *penA* construct, with concentrations of cefixime slightly above the MIC_CFX_ of FA19 (0.004-0.006 µg/ml) for the β3-β4 mutant and up to 0.25 µg/ml for the *penA19*-6M, -7M, and -10M constructs. The next day, colonies were streaked onto fresh GCB plates, and after overnight growth, individual colonies from the streaks were boiled in 1X TE for 5 min, the lysate was centrifuged, and 1-2 µL was used as template for PCR as described below. All transformants were sequence verified before proceeding with MIC experiments.

### Generation of N. gonorrhoeae strains expressing mutant penA sequences

DNA used to transform FA19 was either sequence-verified plasmid constructs or PCR products. For plasmid constructs, sequences were cloned into pUC18us, a vector containing the 10-bp *N. gonorrhoeae* uptake sequence (5’-GCCGTCTGAA-3’). The cloned fragment comprised sequence starting at codon-45 of the *penA* gene and ending at codon-100 of *murE*, the gene downstream of *penA* in the *Neisseria gonorrhoeae* genome. The *penA* gene contained silent restriction sites to split the sequence into 6 modules as previously described [36] (see Supplemental Results and Fig. S1), and the extra *murE* sequence was included to ensure incorporation of mutations close to the 3’-end of the *penA* gene. The template DNA for PCR reactions was either genomic DNA from FA19, H041, and variant strains (extracted using the Promega Wizard Genomic DNA Purification Kit) or from diluted *penA* plasmid DNA. Primers were designed to incorporate specific mutations and comprised ∼15 bases both before and after the desired mutations to promote annealing, with the 5’- and 3’-primers complementary to each other. For certain constructs (i.e. those that conferred small increases in resistance above that for FA19), a silent restriction site was included within the mutation primer to assist in screening of transformants. PCR reactions were carried out with Pfu or Phusion polymerase. Two PCR fragments per mutant were generated: the “upstream” fragment used 5’-*penA*_45aa_US and the 3’-mutation primer, and the “downstream” fragment used the 5’-mutation primer and the 3’-*murE*_31aa primer (Table S4). When the mutations were too distant to incorporate with one set of mutation primers (e.g. the epistatic mutations and the *penA19*-6M and -10M constructs), two or more rounds of overlap extension were carried out. The resulting purified fragments were isolated from agarose gels using Xcluda type D tips, and the full length *penA-murE19* product was amplified by PCR using the outside primers and the upstream and downstream PCR fragments. PCR products were purified using Qiagen QIAquick^®^ PCR Purification Kit and the DNA concentrations determined on a NanoDrop One (Thermo Fisher). These PCR products were used either for ligation into digested plasmid DNA or directly for transformation into *N. gonorrhoeae*. A list of FA19 strains used in this study are shown in Table S5.

### Minimum Inhibitory Concentration (MIC) Evaluations

On the day of the experiment, 25 mL GCB agar plates with varying concentrations of ceftriaxone, cefixime, and penicillin G (2-fold and 1.5-fold dilution series, one plate at each concentration) were poured, allowed to solidify, then dried in a 37 °C incubator for at least 2 hours. The day before, nonpiliated (P-) strains were streaked from frozen stocks onto GC agar plates, and the next day the cells were resuspended in GCB+ broth at an OD_600_ of 0.018 (∼1 x 10^7^ cfu). Five µL of each strain (∼50,000 cfu) were spotted onto each antibiotic plate and incubated for 24 hr. Growth was defined as at least five colonies growing within the spot. MIC experiments were repeated 3 times.

### Analysis of Neisseria species penA loci

PubMLST comprises a curated database of genomic sequences from *N. gonorrhoeae*, *N. meningitidis*, and 40 other *Neisseria* species [40]. From this collection, all unique *penA* genes were aligned using the Locus Explorer plugin on the site. PubMLST has multiple options for calling *penA* loci, including a defined, *N. gonorrhoeae-*only option (listed as “NG_penA”) and an option containing equivalent *penA* loci across all available species within the database (listed as “NEIS1753”). As of this writing, Locus Explorer can align only 2,000 protein sequences at a time, so the 5,274 loci in the NEIS1753 category were aligned in three separate batches, and then manually compared across all three groups.

For searches on the prevalence of the T483S mutation in *Neisseria* species, we used the PubMLST database genome collection tool [40], specifying the species of interest, then searching for the NEIS1753 T483S sequence variation. For searching the other 40 *Neisseria* species for T483S alterations, we excluded both *Neisseria gonorrhoeae* and *Neisseria meningitidis* species from the genome search.

### Protein expression and purification

We previously reported the cloning of a TPase domain construct (designated by the prefix “t”) that we use for structural and biochemical investigations of PBP2 variants [32, 33, 50]. Omission of the N-terminal pedestal domain does not alter second-order rates of acylation for β-lactam antibiotics [50]. A gene construct of *penA19* containing the 10 mutations was synthesized *de novo* (GenScript, Piscataway, NJ) and cloned in-frame into the pMALC2KV vector. This expresses tPBP2 as an N-terminal hexa-histidine–tagged fusion with maltose-binding protein, separated by a tobacco etch virus (TEV) protease site. The tPBP2-6M and -7M variants were then generated by site-directed mutagenesis of *penA19-*10M using the QuikChange Lightning kit (Agilent, Santa Clara, CA). All constructs were confirmed by Sanger sequencing. The resulting plasmids were transformed into *Escherichia coli* BL21 (DE3) cells and proteins expressed and purified as described previously [32]. As a final step, the protein was eluted from a 5 mL HiTrap SP FF cation-exchange column (GE Healthcare, Piscataway, NJ) equilibrated in TG buffer (20 mM Tris-HCl, pH 8.0, 10% glycerol) using a gradient of 0-500 NaCl. Fractions containing the tPBP2 variants were pooled, concentrated, and either used immediately or aliquoted and stored at −80°C.

### Structure determination

Crystals of tPBP2-6M, -7M and -10M were generated by the sitting-drop vapor diffusion method in 96-well plate format, following published protocols (Singh, 2019). Prior to crystallization, proteins were concentrated to 13 mg/ml in the same buffer as eluted from the cation-exchange column. After incubation at 18°C, crystals appeared after 3-4 days with wells containing 32−40% PEG 600 buffered with 0.1 M CHES at pH 9−10. These exhibited a similar plate morphology as crystals as tPBP2^FA19^ and are in the same P2_1_ space group with two molecules in the asymmetric unit. Diffraction data from cryo-cooled crystals were collected at the SER-CAT beamlines at the Advanced Photon Source in Argonne, IL. Data from crystals of tPBP2-6M and tPBP2-7M data were collected at a wavelength of 1.0 Å on an Eiger 16M detector at the ID-22 beamline. 360° of data were collected in 0.25° oscillations at a crystal-to-detector distance of 262 mm (tPBP2-7M) or 280 mm (tPBP2-6M). Data from tPBP2-10M crystals were collected at the BM-22 beamline on a MAR MX300-HS detector. 360° data were collected in 1° increments at a crystal-to-detector distance of 180 mm. After data processing using HKL2000 [51], structures were solved by simple refinement against the tPBP2^FA19^ structure (PDB ID 6P53; [32]) or in the case of tPBP2-10M, by molecular replacement using PHASER [52] using the same starting model. The mutations were introduced into each of the three models by inspecting |Fo|−|Fc| difference electron density maps and followed by iterative cycles of model building and refinement using COOT [53] and REFMAC [54]. Molecule B is used for the structural comparisons since its β3-β4 loop is ordered, compared to it being disordered in molecule A.

### Acylation Rate determination of Minimal Mutant Constructs

The second-order rate constants (*k_2_*/K_s_) of acylation with [^14^C]penicillin G (Moravek, Brea, CA) for the PBP2 variants were determined using purified protein, as described previously [24]. The *k_2_*/K_s_ values for ceftriaxone and cefixime were determined indirectly by determining the concentrations of the cephalosporins that inhibited the binding of [^14^C]penicillin G by 50% [37, 43].

### Quantitative Growth Curves

To generate the FA19 strains used in quantitative growth curves, the full *penA* genes from each minimal mutant construct and FA19 *penA41* were amplified by PCR with 5’-*penA*_45aa_US and 3’-*murE*_31aa primers by colony PCR or genomic DNA. The PCR products were validated by DNA sequencing (GENEWIZ from Azenta Life Sciences), then transformed into the same piliated FA19 stock on the same day. Quantitative growth curves as a measure of fitness were done similarly as described previously [47]. Briefly, after growing overnight on GCB+agar plates, P-strains of *N. gonorrhoeae* were resuspended in GCB+ broth without sodium bicarbonate. The resuspended cultures were used to inoculate 10 mL of GCB+ broth without sodium bicarbonate to a starting OD_600_ of 0.08 in 50 mL tissue culture flasks (BioLite, ThermoFisher). The flasks were grown in a CO_2_ incubator at 180 rpm for a total of 8 hours, with the OD_600_ measured every hour. At the end of the incubation, a sterile loop was dipped in each culture and streaked individually on GCB+ agar plates and assessed for potential culture contamination. Experiments were repeated at least 3 times.

## Acknowledgements

This work was supported by National Institutes of Health awards GM66861 and AI164794 (to C.D. and R.A.N.) and U19 AI113170 (to R.A.N.). Use of the Advanced Photon Source was supported by the U.S. Department of Energy, Office of Science, Office of Basic Energy Sciences, under Contract No. W-31-109-ENG-38. Data were collected at Southeast Regional Collaborative Access Team (SER-CAT) 22-ID and 22-BM beam lines at the Advanced Photon Source, Argonne National Laboratory. Supporting institutions may be found at https://www.ser-cat.org/members.html.

## Notes

### Competing Interest Statement

The authors have declared no competing interest.

## References

1. Unemo M, Seifert HS, Hook EW, 3rd, Hawkes S, Ndowa F, Dillon JR. Gonorrhoea. Nat Rev Dis Primers. 2019;5(1):79. Epub 20191121. doi: 10.1038/s41572-019-0128-6. PMID: 31754194.

2. CDC. Sexually Transmitted Infections Surveillance, 2024 (Provisional)2024. Available from: https://www.cdc.gov/sti-statistics/annual/index.html.

3. Lovett A, Duncan JA. Human Immune Responses and the Natural History of *Neisseria gonorrhoeae* Infection. Front Immunol. 2018;9:3187. Epub 20190219. doi: 10.3389/fimmu.2018.03187. PMID: 30838004.

4. Chang SX, Chen KK, Liu XT, Xia N, Xiong PS, Cai YM. Cross-sectional study of asymptomatic *Neisseria gonorrhoeae* and *Chlamydia trachomatis* infections in sexually transmitted disease related clinics in Shenzhen, China. PLoS One. 2020;15(6):e0234261. Epub 20200609. doi: 10.1371/journal.pone.0234261. PMID: 32516318.

5. Farley TA, Cohen DA, Elkins W. Asymptomatic sexually transmitted diseases: the case for screening. Prev Med. 2003;36(4):502–9. doi: 10.1016/s0091-7435(02)00058-0. PMID: 12649059.

6. Eisinger RW, Erbelding E, Fauci AS. Refocusing Research on Sexually Transmitted Infections. J Infect Dis. 2020;222(9):1432–4. doi: 10.1093/infdis/jiz442. PMID: 31495889.

7. Unemo M, Jensen JS. Antimicrobial-resistant sexually transmitted infections: gonorrhoea and *Mycoplasma genitalium*. Nature reviews Urology. 2017;14(3):139–52. Epub 20170110. doi: 10.1038/nrurol.2016.268. PMID: 28072403.

8. Unemo M, Ross J, Serwin AB, Gomberg M, Cusini M, Jensen JS. 2020 European guideline for the diagnosis and treatment of gonorrhoea in adults. Int J STD AIDS. 2020:956462420949126. Epub 20201029. doi: 10.1177/0956462420949126. PMID: 33121366.

9. Fifer H, Doumith M, Rubinstein L, Mitchell L, Wallis M, Singh S, et al. Ceftriaxone-resistant *Neisseria gonorrhoeae* detected in England, 2015-24: an observational analysis. J Antimicrob Chemother. 2024;79(12):3332–9. doi: 10.1093/jac/dkae369. PMID: 39417254.

10. Lahra MM, Martin I, Demczuk W, Jennison AV, Lee KI, Nakayama SI, et al. Cooperative Recognition of Internationally Disseminated Ceftriaxone-Resistant *Neisseria gonorrhoeae* Strain. Emerg Infect Dis. 2018;24(4):735–40. doi: 10.3201/eid2404.171873. PMID: 29553335.

11. Unemo M, Golparian D, Shafer WM. Challenges with gonorrhea in the era of multi-drug and extensively drug resistance - are we on the right track? Expert review of anti-infective therapy. 2014;12(6):653–6. Epub 2014/04/08. doi: 10.1586/14787210.2014.906902. PMID: 24702589.

12. Unemo M, Del Rio C, Shafer WM. Antimicrobial Resistance Expressed by *Neisseria gonorrhoeae*: A Major Global Public Health Problem in the 21st Century. Microbiology spectrum. 2016;4(3). Epub 2016/06/24. doi: 10.1128/microbiolspec.EI10-0009-2015. PMID: 27337478.

13. Workowski KA, Bachmann LH, Chan PA, Johnston CM, Muzny CA, Park I, et al. Sexually Transmitted Infections Treatment Guidelines, 2021. MMWR Recomm Rep. 2021;70(4):1–187. Epub 20210723. doi: 10.15585/mmwr.rr7004a1. PMID: 34292926.

14. Macheboeuf P, Contreras-Martel C, Job V, Dideberg O, Dessen A. Penicillin binding proteins: key players in bacterial cell cycle and drug resistance processes. FEMS Microbiol Rev. 2006;30(5):673–91. doi: 10.1111/j.1574-6976.2006.00024.x. PMID: 16911039.

15. Sauvage E, Kerff F, Terrak M, Ayala JA, Charlier P. The penicillin-binding proteins: structure and role in peptidoglycan biosynthesis. FEMS Microbiol Rev. 2008;32(2):234–58. Epub 2008/02/13. doi: 10.1111/j.1574-6976.2008.00105.x. PMID: 18266856.

16. Barbour AG. Properties of penicillin-binding proteins in *Neisseria gonorrhoeae*. Antimicrob Agents Chemother. 1981;19(2):316–22.

17. Ropp PA, Nicholas RA. Cloning and characterization of the *ponA* gene encoding penicillin-binding protein 1 from *Neisseria gonorrhoeae* and *Neisseria meningitidis*. J Bacteriol. 1997;179(8):2783–7.

18. Taguchi A, Welsh MA, Marmont LS, Lee W, Sjodt M, Kruse AC, et al. FtsW is a peptidoglycan polymerase that is functional only in complex with its cognate penicillin-binding protein. Nat Microbiol. 2019;4(4):587–94. Epub 20190128. doi: 10.1038/s41564-018-0345-x. PMID: 30692671.

19. Sjodt M, Rohs PDA, Gilman MSA, Erlandson SC, Zheng S, Green AG, et al. Structural coordination of polymerization and crosslinking by a SEDS-bPBP peptidoglycan synthase complex. Nat Microbiol. 2020;5(6):813–20. Epub 20200309. doi: 10.1038/s41564-020-0687-z. PMID: 32152588.

20. Li Y, Gong H, Zhan R, Ouyang S, Park KT, Lutkenhaus J, et al. Genetic analysis of the septal peptidoglycan synthase FtsWI complex supports a conserved activation mechanism for SEDS-bPBP complexes. PLoS Genet. 2021;17(4):e1009366. Epub 20210415. doi: 10.1371/journal.pgen.1009366. PMID: 33857142.

21. Ghuysen J-M, Dive G. Biochemistry of the penicilloyl serine transferases. In: Ghuysen J-M, Hakenbeck R, editors. Bacterial Cell Walls. Amsterdam: Elsevier Science B.V.; 1994. p. 103–29.

22. Ghuysen JM. Serine β-lactamases and penicillin-binding proteins. Annu Rev Microbiol. 1991;45:37–67.

23. Ropp PA, Hu M, Olesky M, Nicholas RA. Mutations in *ponA*, the gene encoding penicillin-binding protein 1, and a novel locus, *penC*, are required for high-level chromosomally mediated penicillin resistance in *Neisseria gonorrhoeae*. Antimicrob Agents Chemother. 2002;46(3):769–77.

24. Powell AJ, Tomberg J, Deacon AM, Nicholas RA, Davies C. Crystal structures of penicillin-binding protein 2 from penicillin-susceptible and -resistant strains of *Neisseria gonorrhoeae* reveal an unexpectedly subtle mechanism for antibiotic resistance. J Biol Chem. 2009;284(2):1202–12. PMID: 18986991.

25. Spratt BG. Hybrid penicillin-binding proteins in penicillin-resistant strains of *Neisseria gonorrhoeae*. Nature. 1988;332(6160):173–6. doi: 10.1038/332173a0. PMID: 3126399.

26. Ameyama S, Onodera S, Takahata M, Minami S, Maki N, Endo K, et al. Mosaic-like structure of penicillin-binding protein 2 Gene (*penA*) in clinical isolates of *Neisseria gonorrhoeae* with reduced susceptibility to cefixime. Antimicrob Agents Chemother. 2002;46(12):3744–9. PMID: 12435671.

27. Tanaka M, Nakayama H, Tunoe H, Egashira T, Kanayama A, Saika T, et al. A remarkable reduction in the susceptibility of *Neisseria gonorrhoeae* isolates to cephems and the selection of antibiotic regimens for the single-dose treatment of gonococcal infection in Japan. J Infect Chemother. 2002;8(1):81–6. Epub 2002/04/17. doi: 10.1007/s101560200011. PMID: 11957125.

28. Lindberg R, Fredlund H, Nicholas RA, Unemo M. *Neisseria gonorrhoeae* isolates with reduced susceptibility to cefixime and ceftriaxone: association with genetic polymorphisms in *penA*, *mtrR*, *porB1b*, and *ponA*. Antimicrob Agents Chemother. 2007;51(6):2117–22.

29. Ito M, Deguchi T, Mizutani KS, Yasuda M, Yokoi S, Ito S, et al. Emergence and spread of *Neisseria gonorrhoeae* clinical isolates harboring mosaic-like structure of penicillin-binding protein 2 in Central Japan. Antimicrob Agents Chemother. 2005;49(1):137–43. doi: 10.1128/AAC.49.1.137-143.2005. PMID: 15616287.

30. Ohnishi M, Golparian D, Shimuta K, Saika T, Hoshina S, Iwasaku K, et al. Is *Neisseria gonorrhoeae* initiating a future era of untreatable gonorrhea?: detailed characterization of the first strain with high-level resistance to ceftriaxone. Antimicrob Agents Chemother. 2011;55(7):3538–45. Epub 2011/05/18. doi: 10.1128/aac.00325-11. PMID: 21576437.

31. Ohnishi M, Saika T, Hoshina S, Iwasaku K, Nakayama S, Watanabe H, et al. Ceftriaxone-resistant *Neisseria gonorrhoeae*, Japan. Emerg Infect Dis. 2011;17(1):148–9. Epub 2011/01/05. PMID: 21192886.

32. Singh A, Tomberg J, Nicholas RA, Davies C. Recognition of the beta-lactam carboxylate triggers acylation of Neisseria gonorrhoeae penicillin-binding protein 2. J Biol Chem. 2019;294(38):14020–32. Epub 20190730. doi: 10.1074/jbc.RA119.009942. PMID: 31362987.

33. Singh A, Turner JM, Tomberg J, Fedarovich A, Unemo M, Nicholas RA, et al. Mutations in penicillin-binding protein 2 from cephalosporin-resistant *Neisseria gonorrhoeae* hinder ceftriaxone acylation by restricting protein dynamics. J Biol Chem. 2020;295(21):7529–43. Epub 20200406. doi: 10.1074/jbc.RA120.012617. PMID: 32253235.

34. Ghuysen JM. Bacterial active-site serine penicillin-interactive proteins and domains: mechanism, structure, and evolution. RevInfectDis. 1988;10:726–32.

35. Takahata S, Senju N, Osaki Y, Yoshida T, Ida T. Amino acid substitutions in mosaic penicillin-binding protein 2 associated with reduced susceptibility to cefixime in clinical isolates of *Neisseria gonorrhoeae*. Antimicrob Agents Chemother. 2006;50(11):3638–45. PMID: 16940068.

36. Tomberg J, Unemo M, Davies C, Nicholas RA. Molecular and structural analysis of mosaic variants of penicillin-binding protein 2 conferring decreased susceptibility to expanded-spectrum cephalosporins in *Neisseria gonorrhoeae*: role of epistatic mutations. Biochemistry. 2010;49(37):8062–70. Epub 2010/08/14. doi: 10.1021/bi101167x. PMID: 20704258.

37. Tomberg J, Unemo M, Ohnishi M, Davies C, Nicholas RA. Identification of amino acids conferring high-level resistance to expanded-spectrum cephalosporins in the *penA* gene from *Neisseria gonorrhoeae* strain H041. Antimicrob Agents Chemother. 2013;57(7):3029–36. Epub 2013/04/17. doi: 10.1128/aac.00093-13. PMID: 23587946.

38. Fenton BA, Tomberg J, Sciandra CA, Nicholas RA, Davies C, Zhou P. Mutations in PBP2 from ceftriaxone-resistant *Neisseria gonorrhoeae* alter the dynamics of the beta3-beta4 loop to favor a low-affinity drug-binding state. J Biol Chem. 2021;297(4):101188. Epub 20210913. doi: 10.1016/j.jbc.2021.101188. PMID: 34529975.

39. Nakayama S, Shimuta K, Furubayashi K, Kawahata T, Unemo M, Ohnishi M. New Ceftriaxone- and Multidrug-Resistant *Neisseria gonorrhoeae* Strain with a Novel Mosaic penA Gene Isolated in Japan. Antimicrob Agents Chemother. 2016;60(7):4339–41. Epub 2016/04/14. doi: 10.1128/aac.00504-16. PMID: 27067334.

40. Jolley KA, Bray JE, Maiden MCJ. Open-access bacterial population genomics: BIGSdb software, the PubMLST.org website and their applications. Wellcome Open Res. 2018;3:124. Epub 20180924. doi: 10.12688/wellcomeopenres.14826.1. PMID: 30345391.

41. Chen M, Shao Y, Luo J, Yuan L, Wang M, Chen M, et al. Penicillin and Cefotaxime Resistance of Quinolone-Resistant *Neisseria meningitidis* Clonal Complex 4821, Shanghai, China, 1965-2020. Emerg Infect Dis. 2023;29(2):341–50. doi: 10.3201/eid2902.221066. PMID: 36692352.

42. Unitt A, Maiden M, Harrison O. Characterizing the diversity and commensal origins of penA mosaicism in the genus *Neisseria*. Microb Genom. 2024;10(2). doi: 10.1099/mgen.0.001209. PMID: 38381035.

43. Frere JM, Nguyen-Disteche M, Coyette J, Joris B. Mode of action: Interaction with the penicillin binding proteins. The chemistry of β-lactams. Glasgow: Chapman & Hall; 1992. p. 148–96.

44. Unemo M, Golparian D, Nicholas R, Ohnishi M, Gallay A, Sednaoui P. High-level cefixime-and ceftriaxone-resistant *N. gonorrhoeae* in France: novel *penA* mosaic allele in a successful international clone causes treatment failure. Antimicrob Agents Chemother. 2011;56(3):1273–80. Epub 2011/12/14. doi: 10.1128/aac.05760-11. PMID: 22155830.

45. Turner JM, Stratton CM, Bala S, Cardenas Alvarez M, Nicholas RA, Davies C. Ureidopenicillins Are Potent Inhibitors of Penicillin-Binding Protein 2 from Multidrug-Resistant *Neisseria gonorrhoeae* H041. ACS infectious diseases. 2024;10(4):1298–311. Epub 20240306. doi: 10.1021/acsinfecdis.3c00713. PMID: 38446051.

46. Zhou K, Chen SC, Yang F, van der Veen S, Yin YP. Impact of the gonococcal FC428 penA allele 60.001 on ceftriaxone resistance and biological fitness. Emerg Microbes Infect. 2020;9(1):1219–29. doi: 10.1080/22221751.2020.1773325. PMID: 32438866.

47. Vincent LR, Kerr SR, Tan Y, Tomberg J, Raterman EL, Dunning Hotopp JC, et al. In Vivo-Selected Compensatory Mutations Restore the Fitness Cost of Mosaic *penA* Alleles That Confer Ceftriaxone Resistance in *Neisseria gonorrhoeae*. mBio. 2018;9(2). Epub 20180403. doi: 10.1128/mBio.01905-17. PMID: 29615507.

48. Guymon LF, Walstad DL, Sparling PF. Cell envelope alterations in antibiotic-sensitive and-resistant strains of *Neisseria gonorrhoeae*. J Bacteriol. 1978;136(1):391–401. doi: 10.1128/jb.136.1.391-401.1978. PMID: 101519.

49. Kellogg DS, Peacock WL, Deacon WE, Browh L, Perkle CI. *Neisseria gonorrhoeae*. I. Virulence genetically linked to colonial variation. J Bacteriol. 1963;85:1274–9.

50. Fedarovich A, Cook E, Tomberg J, Nicholas RA, Davies C. Structural effect of the Asp345a insertion in penicillin-binding protein 2 from penicillin-resistant strains of *Neisseria gonorrhoeae*. Biochemistry. 2014;53(48):7596–603. Epub 2014/11/19. doi: 10.1021/bi5011317. PMID: 25403720.

51. Otwinowski Z, Minor W. Processing of X-ray diffraction data collected in oscillation mode. Methods Enzymol. 1997;276:307–26.

52. McCoy AJ, Grosse-Kunstleve RW, Adams PD, Winn MD, Storoni LC, Read RJ. Phaser crystallographic software. J Appl Crystallogr. 2007;40(Pt 4):658-74. Epub 20070713. doi: 10.1107/S0021889807021206. PMID: 19461840.

53. Emsley P, Lohkamp B, Scott WG, Cowtan K. Features and development of Coot. Acta crystallographica Section D, Biological crystallography. 2010;66(Pt 4):486–501. Epub 20100324. doi: 10.1107/S0907444910007493. PMID: 20383002.

54. Murshudov GN, Vagin AA, Dodson EJ. Refinement of macromolecular structures by the maximum-likelihood method. Acta crystallographica Section D, Biological crystallography. 1997;53(Pt 3):240–55. doi: 10.1107/S0907444996012255. PMID: 15299926.

